# AI-Guided Precision in Antibody Humanization: Structural Modeling to Minimize Immunogenicity and Preserve Efficacy

**DOI:** 10.1101/2025.03.09.641110

**Authors:** Haiying Wang, Chunqiu Xia, Yupeng Zhu, Yali Wang, Xinxin Chu, Xuemin Pan, Hao Zhou, Zhen Lu, Patrick Doonan, Yi Li, Yunfei Long, Quan Yu

## Abstract

Antibody humanization is an essential part of converting animal-derived antibodies into clinical candidates however conventional CDR grafting techniques often faces challenges, especially when essential antigen-binding sites extend beyond CDR regions. Additionally, the CDR grafting process tends to retain undesired non-human elements within the CDR regions. In this study, we overcame these challenges by using our proprietary AI-predicted antigen-antibody complex algorithm to effectively humanize a murine antibody. During our AI-powered humanization process, in contrast to traditional CDR grafting techniques, only the crucial paratopes identified through our AI-predicted complex structure were precision grafted onto a select human germline. This paratope grafting technique significantly reduced the antibodies immunogenicity risk while maintaining bioactivity. Moreover, all AI-guided deep humanized antibody variants demonstrated favorable developability, including thermal stability, poly-reactivity, and hydrophobicity. These results not only advance antibody humanization techniques but also demonstrate the power of AI in expediting antibody engineering and derisking clinical research.

## Introduction

Therapeutic antibodies have revolutionized drug applications, emerging as pivotal treatments for diverse diseases. As of 2024, over a hundred therapeutic antibodies have been approved, with dozens of others in late-stage clinical trials (Mullard 2021; Crescioli et al. 2025; Mullard 2021). The growing focus on therapeutic monoclonal antibodies and their derivatives, including Fc-fusion proteins, nanobodies, bispecific antibodies, antibody fragments, and antibody-drug conjugates, highlights their importance in medical research (Kaplon et al. 2023). The initial approval of muromonab-CD3 (Todd and Brogden 1989; Wilde and Goa 1996), a murine IgG2a antibody, exposed the immunogenicity issues of animal-derived antibodies, posing challenges in reduced half-lives and safety profiles(Lu et al. 2020). This necessitated the development of humanized antibodies with minimal immunogenicity risk(Safdari et al. 2013; Ahmadzadeh et al. 2014; Waldmann 2019).

The initial attempts to reduce immunogenicity risk entailed substituting the non-human constant regions of the antibody with human sequences to generate chimeric antibodies. This approach evolved with the advent of Complementary-Determining Region (CDR) grafting (Jones et al. 1986), which became the primary humanization strategy (Safdari et al. 2013). However, this traditional technique carries with it risk of failure due to the absence of structural insights into antigen-antibody complexes and often an assumption of an unacceptable decrease in affinity post-humanization.

Therefore, we proposed a novel structure-based humanization strategy that is facilitated by our proprietary complex structure prediction algorithm, named XtalFold^®^. This deep learning attention-based algorithm utilizes multiple sequence alignment (MSA) and homologous templates of the query protein sequences as inputs and outputs the Cartesian coordinates of the structures in complex.

This study leveraged XtalFold^®^ to navigate the humanization of an antibody where traditional approaches had failed due to the critical interaction residue located in a framework region. Our XtalFold^®^ model identified crucial residues within both the CDR and framework regions involved in antigen binding. This allowed us to significantly reduce immunogenicity risk while preserving bioactivity by selectively grafting key paratope residues onto a human framework. This achievement underscores the delicate balance between maintaining affinity and reducing immunogenicity in antibody humanization. Additionally, it exemplifies the transformative impact of AI in antibody engineering and the potential for AI to improve clinical outcomes.

## Results

### High Resolution Computational Prediction of Antigen-Antibody Complex Structure

ALX07 was isolated by traditional mouse hybridoma technology and exhibited comparable bioactivity to the benchmark pembrolizumab analogue (Garon et al. 2015; Kwok et al. 2016; Robert et al. 2015) when expressed as an IgG4 chimeric antibody. To reduce potential immunogenicity risk, humanization was performed using two rounds of CDR grafting which included classical vernier zone back mutations in an effort to retain binding. Multiple humanized variants were evaluated, however, none of the clones had comparable affinity to the parental antibody (Supplementary Figure 1). Considering these results, we suspected that the key antigen-binding residues of ALX07 might extend beyond the CDR regions.

To investigate this suspicion, the complex structure was predicted using our proprietary AI-powered complex modeling algorithm XtalFold^®^. The antibody sequence of ALX07 and the sequence of Programmed cell death protein 1 (PD-1) (Parry et al. 2005; Keir et al. 2008; Topalian, Drake, and Pardoll 2015) were used to generate a high-resolution complex structure without docking or homology modeling. The XtalFold^®^ predicted PD-1 structure demonstrated an RMSD of 0.436Å when aligned with the reference structure (PDB ID: 4ZQK) giving us confidence that we had an accurate structure to guide humanization (Zak et al. 2015). Additionally, XtalFold^®^’s antibody structure model showed a high degree of consistency across all regions when cross-referenced to predictions made by other algorithms, except in CDR3 which is a known challenge for other algorithms (Supplementary Table 1).

**Table 1:** Humanized ALX07 Binding affinity with human PD-1.

| Antibody | $k_a$ (1/Ms) | $k_d$ (1/s) | KD (M) | Back Mutation* |
| --- | --- | --- | --- | --- |
| ALX07xh-05 | 1.59E+05 | 2.22E-02 | 1.40E-07 | - |
| ALX07xh-06 | 4.45E+05 | 8.93E-04 | 2.01E-09 | S67Y |
| ALX07xh-07 | 4.21E+05 | 6.18E-04 | 1.47E-09 | S67Y Y36F Y87F |
| ALX07xh-37 | 1.02E+05 | 1.26E-02 | 1.23E-07 | Y36F Y87F |
| ALX07 | 3.48E+05 | 4.55E-04 | 1.31E-09 | - |

\*Back Mutation: human framework region residues back mutated to murine residues

As speculated, XtalFold^®^’s complex structure prediction revealed an atypical binding pose of ALX07, notably within the light chain (Figure 1A and 1B). Interactions within the heavy chain are primarily localized around HCDR3, as expected. Specifically, Y106 and D108 of HCDR3 interact with T53 and Q52 of PD-1, while N103 of HCDR3 forms two hydrogen bonds with D69 and D54 of PD-1. Notably, within the light chain, each CDR region actively participates in binding. In LCDR1, N31 and D32 interact with antigen residue K55, while D91 and Y92 in LCDR3 engage with Q65 and D62 of the antigen. Although LCDR2 lacks hydrogen bond interactions, the proximity of residue F52 to the β-sheet residues I103, Q110, and I111 of the antigen gives rise to Van der Waals forces. Proximal to the CDRs of the light chain, Y67 in LFR3 interacts with L105 and V41 of PD-1 given their close spatial arrangement (Figure 1C & 1D). Part of the antigen:antibody interface results in the formation of a hydrophobic core formed from the antigen’s three parallel β-sheets including residues V41, I103, L105, A109, and I111 in conjunction with F52 and Y67 from the antibody light chain (Figure 1E).

**Figure 1.**
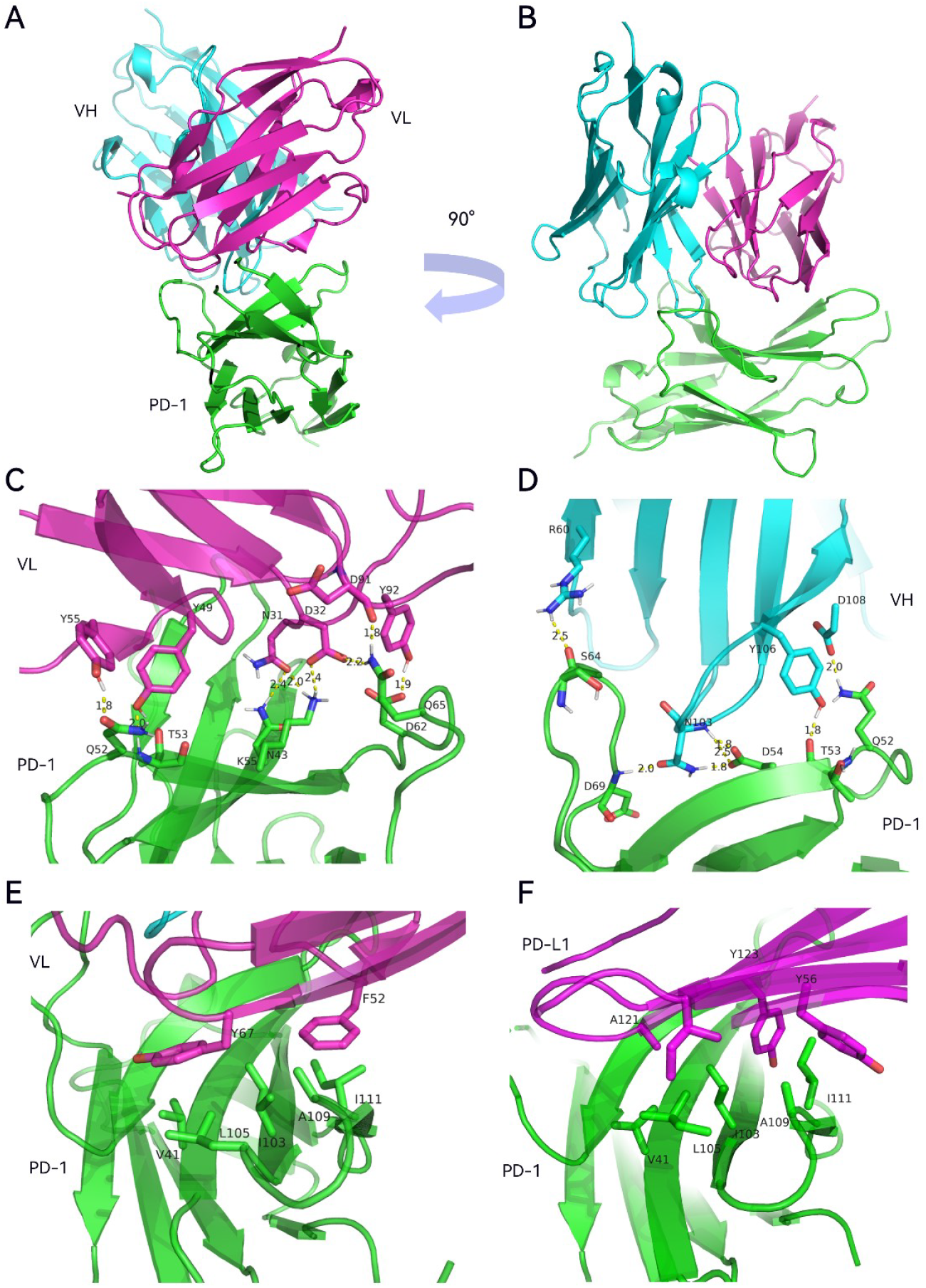
XtalFold^®^ modeling structure illustrates ALX07’s atypical binding pose with human PD-1. In (A) to (E), cyan, purple and green represent VH, VL and human PD-1 extracellular domain respectively. In F, green and purple represent PD-1 and PD-L1 extracellular domain respectively. (A) (B)Predicted structure of ALX07 with PD1 with 90□ rotated view (ribbon backbone). (C). Interactions between VL region and PD-1, side chains visible for residues distances calculated in Angstroms. (D) Interactions between VH region and PD-1. Side chains visible and distance calculated in Angstroms. E. Critical hydrophobic interaction of VL region and PD-1. (F) Hydrophobic interaction of PD-L1 and PD-1 (PDB ID: 4ZQK); PD-1 at similar orientation to E

Our complex structure model reveals that the light chain of ALX07 plays a pivotal role in binding interactions with PD-1, surpassing that of the heavy chain. Interestingly, inspection of the binding sites of PD-1 and PD-L1 (Zak et al. 2015) shows a similar formation of a hydrophobic core in analogous positions to that of our XtalFold^®^ predicted ALX07/PD-1 complex structure (Figure 1F). The XtalFold^®^ structure model of the antigen-antibody complex provided crucial guidance during the re-designing of humanized variants of ALX07.

### AI Modeling Complex Structure Yields Insight into Role of Y67 in Antigen Binding and Rescues Humanization of ALX07

In light of the structural information gleaned from the XtalFold^®^ complex, we initiated a new humanization by grafting the CDRs of ALX07 into human scaffold sequences derived from IGHV 2-70 and IGKV 4-1, which showed high sequence similarity to murine ALX07 (Figure 2A and 2B). The AI-powered complex structure predicted several crucial framework residues to be essential for maintaining CDR conformation and preserving antigen binding. Therefore, these residues were selectively reverted to their murine counterparts as back mutations. Following humanization, the direct CDR-grafted variant (ALX07xh-05) exhibited a significant decrease in affinity (KD 1.40E-07 M, Figure 3C, Table 1). However, reintroducing the S67Y mutation in the light chain alone was sufficient to restore antigen affinity (KD 2.01E-09 M, compared to 1.31E-09 M for the murine antibody, Figure3A and 3E). Other back mutation variants like ALX07xh-37 that lacked S67Y of the light chain failed to recover affinity levels (KD 1.23E-07 M, Figure 3D, Table 1). This finding supports the predictions of the AI complex structure model and emphasizes the critical function of Y67 in the VL framework of murine ALX07 in antigen binding.

**Figure 2.**
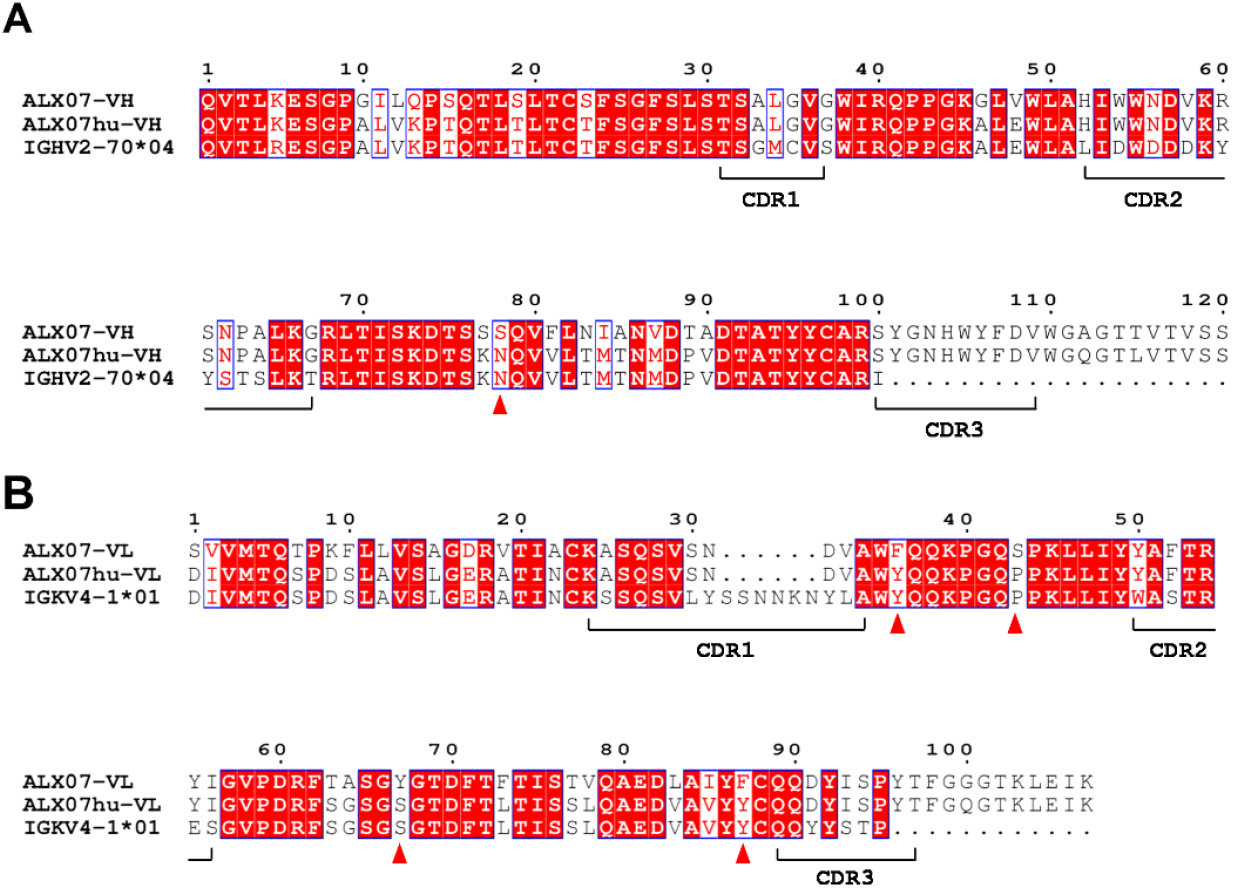
Alignment of CDR-Grafted Humanized Sequences with Murine and Germline Sequences. (A) VH sequence alignment and (B) VL sequence alignment, comparing the murine ALX07 sequence (top row), the humanized sequence (middle row), and the germline sequence (bottom row). Red triangles mark the positions of back mutations introduced to preserve antigen binding and structural integrity.

**Figure 3.**
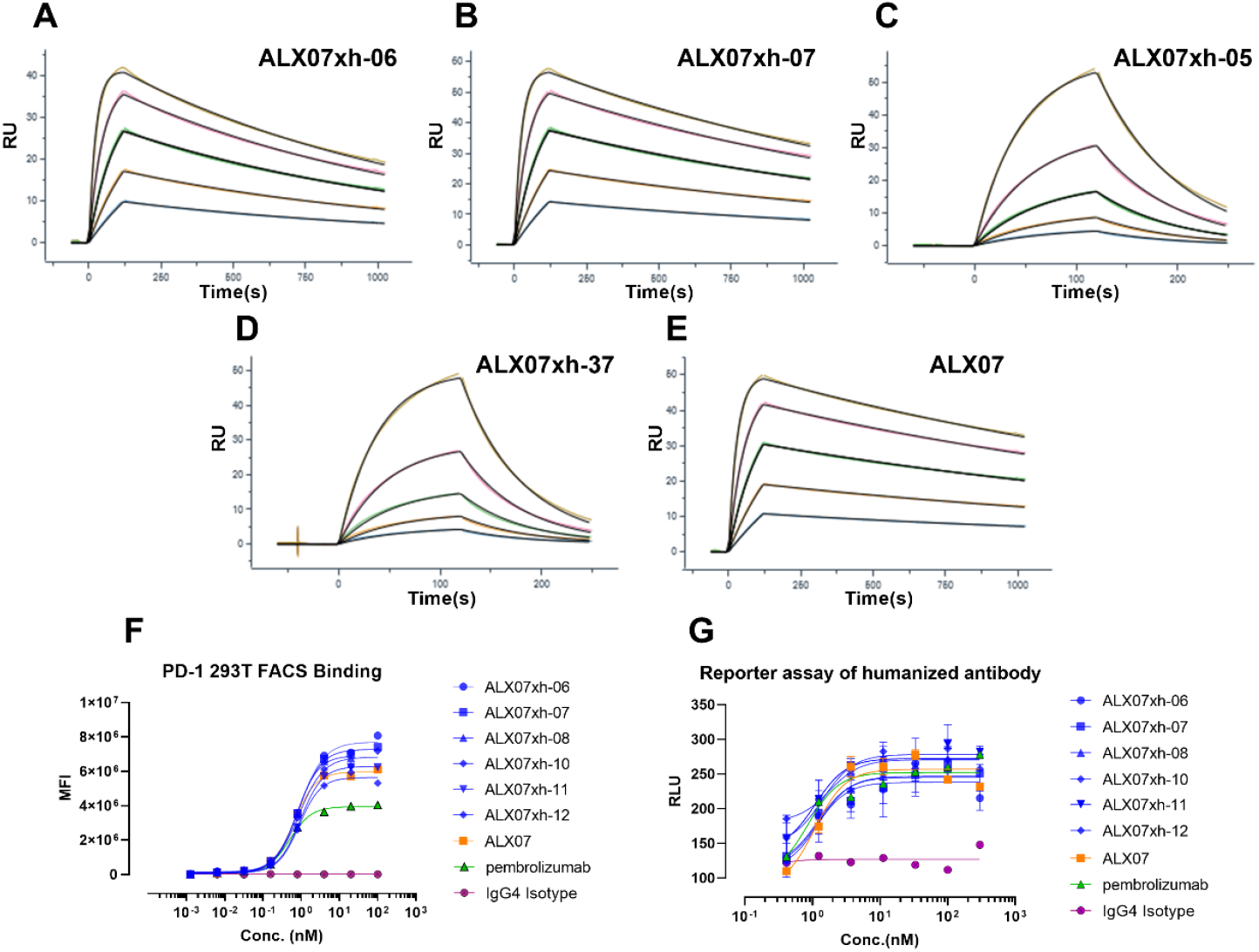
Bioactivity assessment of CDR-graft humanization variants. (A)-(E). Surface plasmon resonance (SPR) analysis of binding affinity of ALX07 variants. (F) Binding of ALX07 variants binding with CHO-PD1 cells; (G) NFAT-luciferase reporter assay results. In panels (F) and (G), blue, orange, green, purple represent humanized variants, chimeric ALX07, a benchmark antibody, and an isotype control, respectively.

Interestingly, Y67 is located in framework 2, distant from any CDR loops, and would typically be replaced by a humanized residue in the absence of antigen-antibody structure information, leading to a loss of bioactivity. By utilizing AI-predictive insights, Y67 was found to be a critical amino acid. Subsequent cellular assays confirmed that the humanized ALX07 with the Y67 backmutation retained its PD-1 binding and blocking efficacy, comparable to the parental murine counterpart (Figure 3F and 3G).

Considering the significance of this framework residue, we examined the amino acid distribution at this locus within the human and murine germlines (Figure 4A), with a focus on the exclusivity of Y67 as a key back mutation. Our analysis revealed that tyrosine is a relatively rare residue in both species. This discovery raised concerns about the immunogenicity of the CDR-grafted molecules containing the essential Y67 back mutation and justified further investigation.

**Figure 4.**
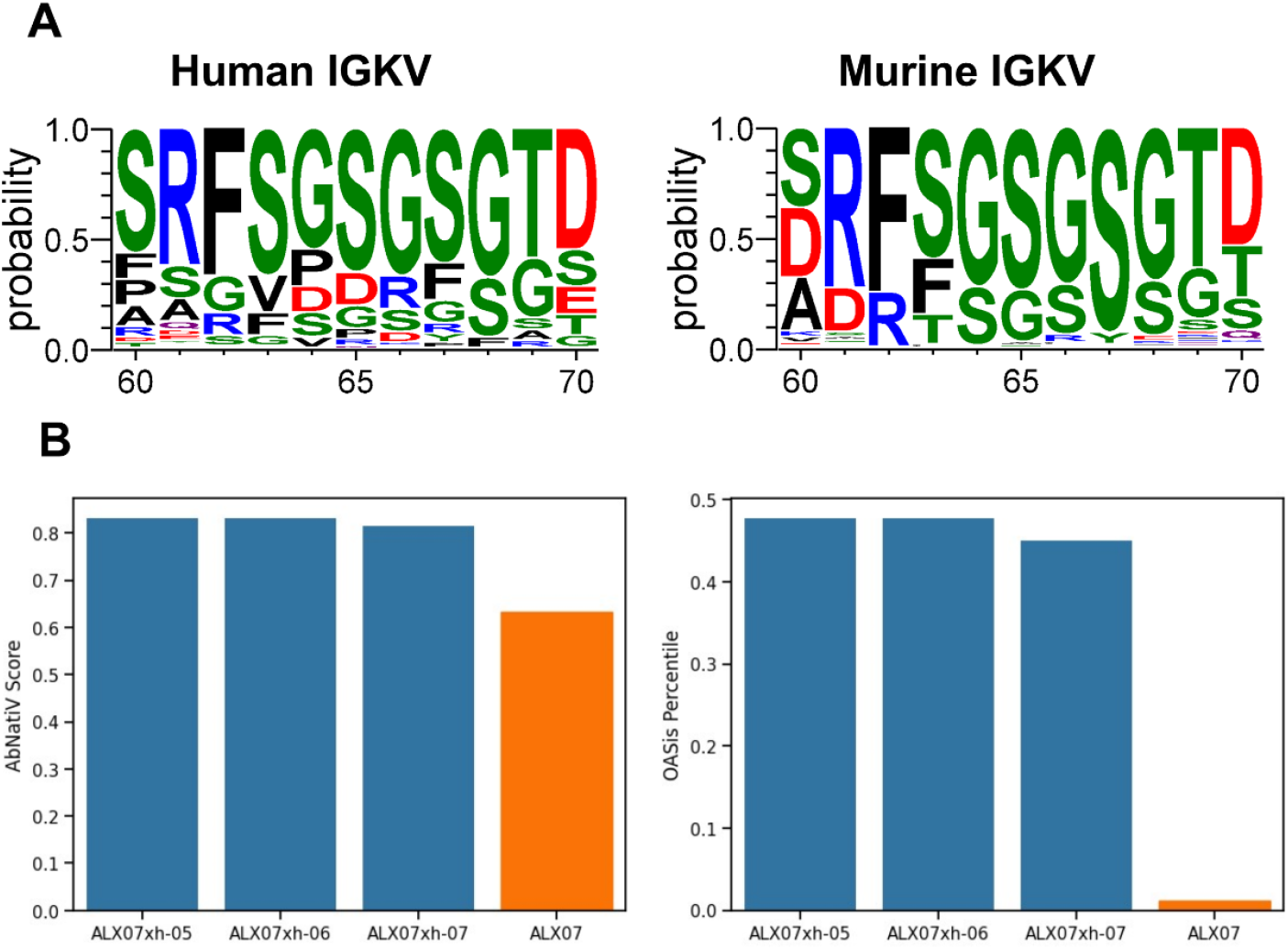
(A). Natural diversity in human and murine IGKV in regions 60 to 70 for each amino acid. Left: human germline; right: murine germline. (B) Humanness evaluation of mAb031 by two algorithms. Left: AbNatiV; right: OASis percentile.

### Enhanced Humanness Level through AI-guided Paratope Grafting

After reviving affinity by inclusion of the essential framework residue Y67, we strived to alleviate immunogenicity risk as predicted by the humanness level of the antibody variants. Using the AbNatiV (Ramon et al. 2024) and OASis (Prihoda et al. 2022) humanness scoring algorithms as a surrogate of immunogenicity risk, we quantified the humanization levels for the murine sequence and compared them with humanized variants, both with and without the essential Y67 mutation.

The OASis percentile score reflects the natural occurrence of peptides formed by an antibody, and its results indicated a remarkably high immunogenicity risk in the chimeric version of ALX07. Expectedly, the humanized versions showed a significant improvement in humanness score relative to the chimeric antibody. Importantly, the Y67 mutation present in variants ALX07xh-06 and ALX07xh-07 did not demonstrate an increase in immunogenicity risk relative to clone ALX07xh-05 which lacked Y67 (Figure 4B).

We wanted to leverage the structural information from XtalFold^®^ in an effort to further reduce the immunogenicity risk of the humanized antibodies. Therefore, we incorporated our AI-powered complex structure model into an innovative humanization protocol introduced here as “paratope grafting.” This method is facilitated by predicted structure and involves grafting the antigen-interacting residues (paratope) (Jerne 1974; Izadinia, Sadeghi, and Ebadzadeh 2009) from a murine antibody onto a human germline, along with the residues necessary for maintaining the paratope’s conformation. While similar to the previously described SDR grafting technique (Padlan, Abergel, and Tipper 1995; Kashmiri et al. 2005),”paratope grafting”utilizes complex structural information and places greater emphasis on paratope conformation. This method not only improves humanness by minimizing the number of murine residues grafted but also maintains affinity by retaining all essential residues.

The paratope grafting humanization strategy involves first selecting an appropriate germline as the accepting framework, followed by identifying key paratope residues and assessing the influence of neighboring residues. The key paratope residues are identified using the XtalFold^®^ complex structure and subsequently grafted onto the chosen germline, after which “back mutations” of crucial neighboring residues on CDRs are introduced to strike a balance between maintaining affinity and achieving the highest level of humanization. This process ultimately yields a maximally human molecule that is only efficiently accomplished due to the sequence-based, AI-predicted complex structure.

As previously noted, the human framework IGKV4-1 was chosen for the light chain, while IGHV2-70 was selected for the heavy chain. Our AI complex structure of chimeric ALX07 and PD-1 revealed 17 amino acids within the CDRs as key paratope residues, comprising only 26% of the total CDR length (Kabat definition). Highlighting the significant amount of parental CDR residues that are unnecessarily included during traditional humanization. Further analysis identified 48 neighboring residues interacting with these key paratope residues. In concert, these supporting residues, which included Y67 as previously identified, support the key contact residues, and are recognized as critical to reformulate the paratope structure in the humanized antibody.

Initial in silico designs included all identified key paratope residues of CDR1 and CDR2 along with the entire murine CDR3. Additionally, a thorough analysis of neighboring residues was conducted to identify potential back mutation sites that might impact CDR loop confirmation. The back mutation candidate residues were evaluated by in-depth structural analysis to assess whether a human residue would deform the conformation of the paratope. If a possible deleterious effect was predicted, the mouse residue was included as a back mutation candidate otherwise, the human residue was retained in an effort to maximize humanness. Based on the potential impact on paratope conformation, various combinations of back mutations were designed. This AI structure-based method significantly reduces resources and improves the likelihood of success at minimizing non-human content in candidate antibodies.

From the entire set of paratope-neighboring residues evaluated (Supplementary Table 2), the light chain contained three neighboring residues that differed between the murine and human sequences (K30, V33 and A25). As shown in Figure 1C and Figure 5A, light chain residues N31 and D32 are both involved in antigen interaction. The human K30, which corresponds to a serine in the murine sequence, was predicted to have the greatest influence on antigen binding. K30 could potentially affect antigen binding in two ways: first, through steric hindrance due to the larger side chain of lysine compared to serine; second, by forming a cis salt bridge with D32, which could reduce the interaction between D32 and the antigen thereby altering the paratope’s electrostatic environment (Figure 5B). In contrast, the other two paratope-neighboring sites, S25 and L33 have similar side chain size as their murine counterparts (A25 and V33, respectively) and inward orientations suggesting a minimal impact (Figure 5B).

**Table 2.** Design of paratope graft.

| Chain | V gene germline | Adjacent residue back mutated number | Back mutation |
| --- | --- | --- | --- |
| VH-70A | IGHV2-70 | 0 | - |
| VH-70B | IGHV2-70 | 5 | S37G R52H D54W S62N T63P |
| VH-70C | IGHV2-70 | 7 | M34L R35G S37G R52H D54W S62N T63P |
| VH-70D | IGHV2-70 | 9 | M34L R35G S37G R52H D54W S62N T63P S64A T67G |
| VL-1A | IGKV4-1 | 1 | K30S |
| VL-1B | IGKV4-1 | 2 | K30S L33V |
| VL-1C | IGKV4-1 | 3 | K30S L33V S25A |

*\*VH-70D is the same heavy chain used in CDR-graft design.*

**Figure 5.**
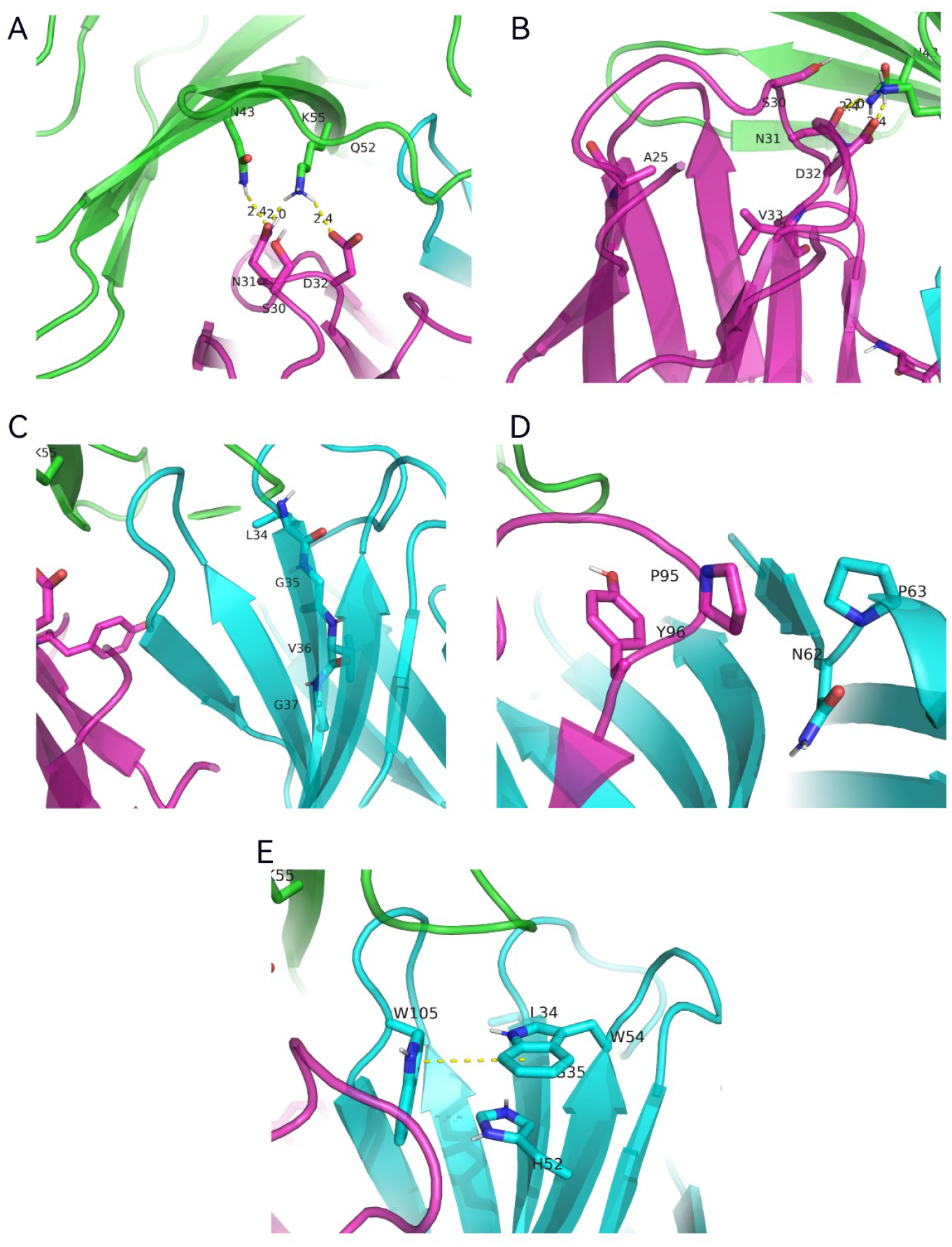
XtalFold^®^ predicted complex guided the design of paratope-graft. On (A) to (E), cyan, purple and green represent VH, VL and human PD-1 extracellular domain, respectively.

While in the heavy chain, the IGHV2-70 germline exhibited seven non-parental sites in paratope-neighboring residues: L34, G35, G37, R52, D54, N62, P63 (Supplementary Table 2). Murine W54 in HCDR2 interacts with W105 in HCDR3, making it a critical site supporting a key contact residue (Figure 5C). Therefore, D54 in the human germline could disrupt this interaction and significantly affect affinity by disturbing the HCDR3 conformation. Similarly, human R52 is located near W105 and may alter HCDR3 conformation due to steric hindrance from the arginine side chain. While murine P63 was shown to be distant from the antigen, it remained a backmutation candidate because of the structural rigidity of proline and its position in the turn which could influence HCDR2 backbone conformation (Figure 5D). Additionally, N62, located in proximity to LCDR3, was predicted to play a role in supporting the LCDR3 structure therefore it was included in the variant designs. Finally, HCDR1, contained three murine residues L34, G35, and G37 that formed an antiparallel β-sheet with HCDR3, which could potentially influence HCDR3 loop conformation (Figure 5E).

Guided by the AI complex structure analysis above, we designed three paratope-grafted light chains and three paratope-grafted heavy chains. Additionally, the heavy chain from the conventional CDR-graft method was incorporated in the designs as VH-70D (Table 2). All heavy and light chain combinations were expressed in the same IgG4 format as previously described. A total of 12 variants were analyzed for affinity by surface plasmon resonance (SPR). As a result, six molecules showed no binding signal and six maintained comparable affinity (Table 3, Figure 6D)

**Table 3.** Affinity of paratope graft variants.

| name | VH chain | VL chain | ka (1/Ms) | kd (1/s) | KD (M) |
| --- | --- | --- | --- | --- | --- |
| ALX07xh-25 | VH-70A | VL-1A | — | — | — |
| ALX07xh-26 | VH-70A | VL-1B | — | — | — |
| ALX07xh-27 | VH-70A | VL-1C | — | — | — |
| ALX07xh-28 | VH-70B | VL-1A | — | — | — |
| ALX07xh-29 | VH-70B | VL-1B | — | — | — |
| ALX07xh-30 | VH-70B | VL-1C | — | — | — |
| ALX07xh-31 | VH-70C | VL-1A | 3.43E+05 | 8.83E-04 | 2.57E-09 |
| ALX07xh-32 | VH-70C | VL-1B | 3.46E+05 | 9.54E-04 | 2.76E-09 |
| ALX07xh-33 | VH-70C | VL-1C | 3.51E+05 | 9.56E-04 | 2.72E-09 |
| ALX07xh-34 | VH-70D | VL-1A | 3.16E+05 | 8.59E-04 | 2.71E-09 |
| ALX07xh-35 | VH-70D | VL-1B | 3.59E+05 | 8.93E-04 | 2.49E-09 |
| ALX07xh-36 | VH-70D | VL-1C | 3.60E+05 | 9.02E-04 | 2.51E-09 |
| ALX07 | parental | parental | 3.12E+05 | 5.68E-04 | 1.82E-09 |

**Figure 6.**
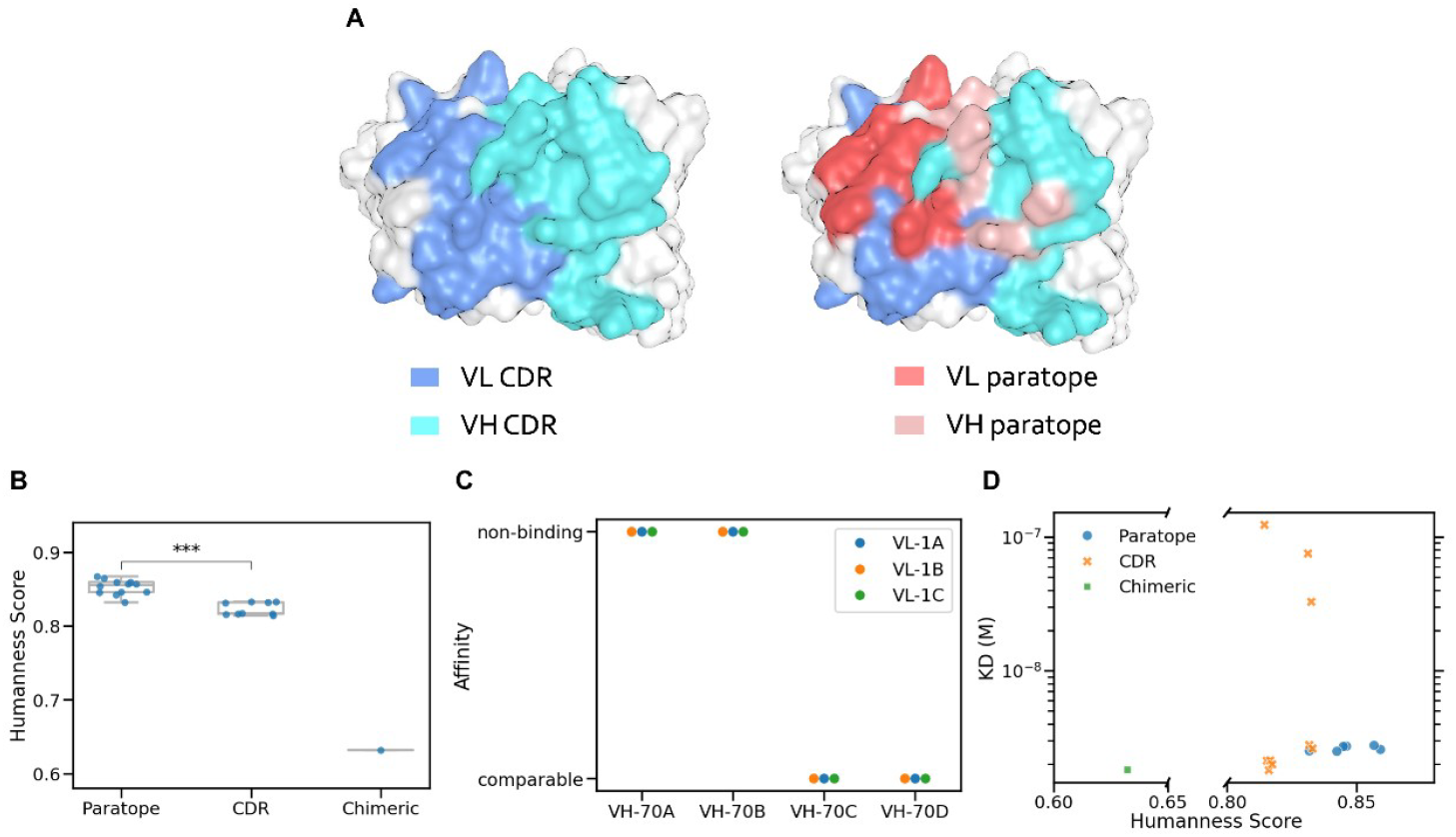
XtalFold^®^ modeling-guided paratope graft humanization design resulted in variants with an improved balance between humanness and bioactivity. (A) CDR and paratope region from the complex model. (B) AbNatiV scores are grouped by design methods. ***p < 0.001 (Student’s t-test). (C) Affinity results of paratope graft variants, grouped by different variable region chains. (D) Scatterplot analysis reveals affinity and humanness levels. Affinity was measured by SPR, and humanness is represented by the AbNatiV score. Each point represents a ALX07 variant. Non-binding variants are not displayed in the figure.

The consistency between the structural observation and experimental results further supports the reliability of the AI-predicted structure. Following the introduction of back mutations at two sites in HCDR1 (M34L and R35G), the non-binder variant ALX07-xh28 recovered its affinity as AL07-xh31. Given that L34 and G35 are buried (with residue accessible surface area percentages of 1.49% and 0%, respectively), it is unlikely that they interact directly with the antigen; instead, they likely affect affinity by influencing the conformation of HCDR3. In the light chain, consistent with the initial analysis, a single back mutation, K30S, was sufficient to restore affinity to that of the parental antibody.

As an effort to predict immunogenicity risk, the humanization levels of the variants were quantified using two algorithms: AbNatiV and OASis percentile. These two algorithms showed a strong correlation in our molecules (Supplementary Figure 2), so we used the AbNatiV score to represent humanness for future analysis. Among all the variants, 7 out of 11 showed improved humanness but reduced affinity. Two variants, ALX07xh-31 and ALX07xh-34 maintained their affinity while achieving notably high humanness (Figure 6D, highlighted with red circle). Regardless of affinity, the paratope-grafted humanization variants demonstrated significantly higher humanness score compared to conventional CDR-grafted humanization (p-value < 0.001, Figure 6B).

Variants ALX07xh-31 and ALX07xh-34, which demonstrated high affinity and high humanness score, were selected for further cell binding and in vitro activity evaluation. These results demonstrate that the paratope grafting candidates exhibit bioactivity comparable to that of chimeric antibody and conventional humanized molecules derived from CDR grafting (Supplementary Figure 5).

### Super-humanized Antibody is Highly Developable

In evaluating our paratope-grafted humanized variants and the conventional humanized variants, we conducted a comprehensive analysis of their biophysical characteristics, including thermal stability, self-association, polyspecificity, and hydrophobic interactions (Figure 7). The biophysical characteritics of these molecules showed minimal variability and did not raise any concerns reagarding developability. Their thermal stability, indicated by melting temperatures (T_m_2, Fabs Tm), were around 80°C. The DNA and BVP binding scores, which reflect antibody polyspecificity, along with AC-SINS measurement for self-association, positioned our molecules near negative controls, indicating satisfactory performance. The Surface-Mounted Adjustable Component (SMAC) analysis, which assesses molecular hydrophobicity, revealed that shorter retention times, as observed in our antibodies compared to pembrolizumab, correlated with weaker hydrophobic interactions. Importantly, the paratope-grafted variants exhibited no devevelopability issues when compared with conventional humanization across all these properties.

**Figure 7.**
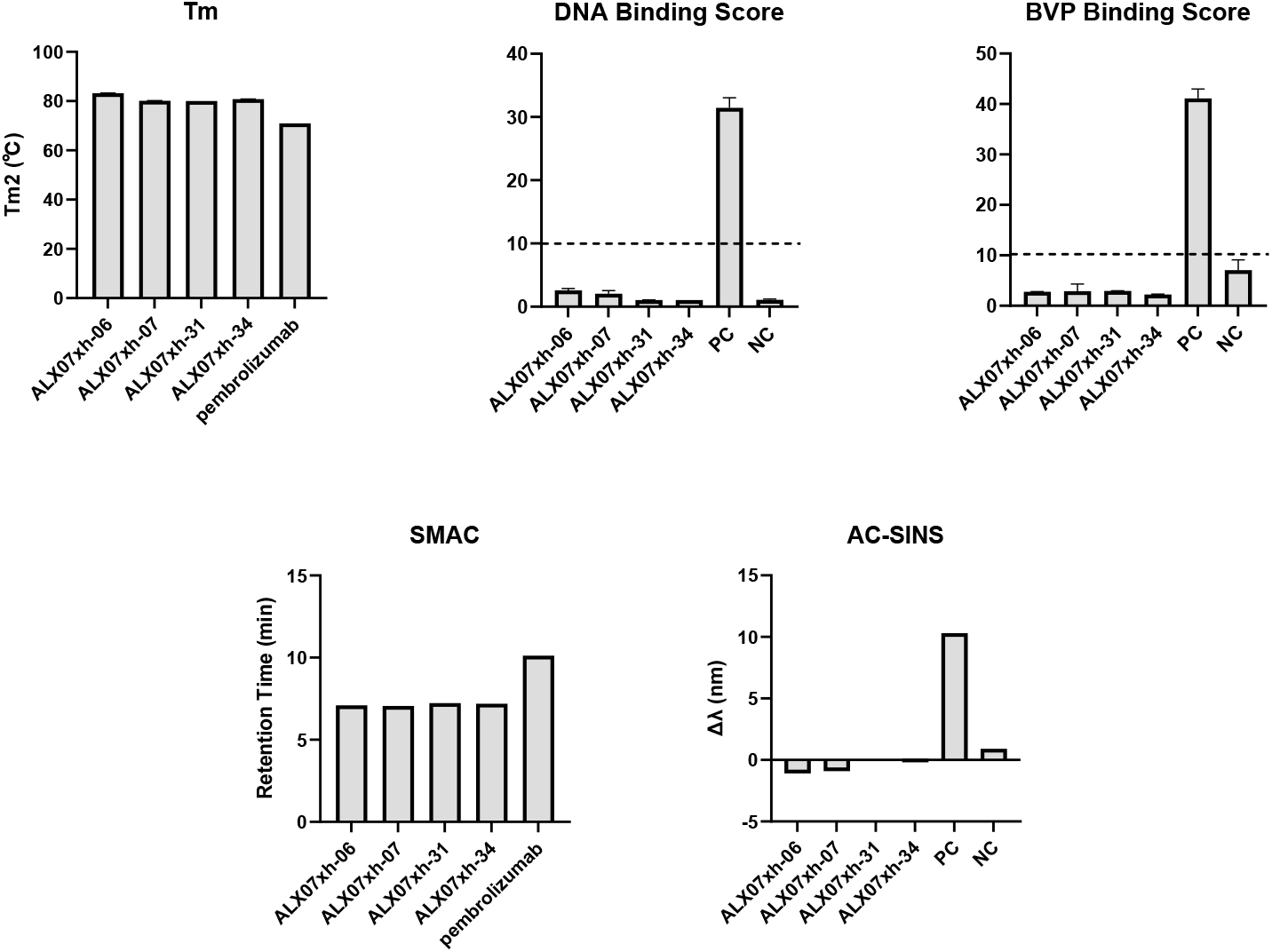
ALX07 humanization variants show favorable biophysical properties in early developability assessments. (A) Tm2 measured by DSF. (B) ELISA binding to DNA. (C) ELISA binding to BVP. (D) Hydrophobicity evaluation by SMAC. (E) self-association measured by AC-SINS. The experiment details refer to the methods.

## Discussion

In recent years, the rapid advancement of computational and artificial intelligence technologies has significantly impacted the biomedical field, particularly enhancing antibody drug development. Our study here explores the application of AI-driven protein complex modeling in the context of antibody humanization.

Although conventional CDR grafting is generally effective, it comes with certain limitations. Traditional methods are both time-consuming and resource-intensive, typically requiring at least three weeks per humanization cycle, and demanding additional iterations if initial attempts prove unsuccessful. Furthermore, this method encounters challenges if the paratope extend s beyond the CDRs. Such limitations were evident in our initial attempts to humanize the murine antibody ALX07, where over 30 humanized molecules were generated but none retained bioactivity.

Utilizing our proprietary AI antibody/antigen complex model XtalFold^®^, we identified a crucial antigen-binding region in the framework area of our PD-1 binder, ALX07. This framework region it typically overlooked during a conventional humanization method that lacks experimental protein structure data. Guided by our AI complex model, we found that just a single back mutation at a key light chain framework site, Y67, following direct CDR grafting onto a human scaffold, was sufficient to maintain bioactivity. This site has been highlighted in previous work, where Y67 was identified as a significant back mutation following the resolution of the antigen-antibody complex structure (Li et al. 2023). This underscores the value of early sequence-based complex structure modeling to achieve what traditionally required high-resolution structure obtained from Cryo-EM or X-ray crystallography.

Reducing antibody immunogenicity risk is the core objective of humanization (Hwang and Foote 2005; Presta 2006). Molecules humanized using CDR grafting result in non-human segments within the CDRs and at the junction of CDRs and FR regions, that drives immunogenicity risk. To further minimize immunogenicity risk, we moved beyond conventional CDR grafting, by grafting only the residues directly interacting with the antigen, along with their supporting residues. Drawing inspiration from previous studies on specificity-determining residues (SDRs) grafting(Padlan, Abergel, and Tipper 1995; Kashmiri et al. 2005), we proposed a concept of ‘paratope grafting.’ This approach utilizes AI predicted complex structure information to identify and graft active residues while also considering adjacent residues that structurally support these paratopes. By transplanting this collective paratope on human frameworks, we were able to maximize the humanness level while preserving bioactivity.

Our findings expand the application of protein complex modeling in antibody engineering. To our knowledge, no published studies have yet utilized AI-predicted antigen-antibody complexes to guide humanization, especially in designing molecules with significantly reduced immunogenicity risk. This study exemplifies the integration of cutting-edge research in protein structure prediction, bioinformatics, and structure-guided protein engineering, highlighting the synergy of interdisciplinary collaboration, a trend likely to shape future advancements. We are optimistic that AI technology will continue to play a pivotal role in protein engineering across a wide range of applications.

## Methods

### Structure Model and inspection

The parental mouse ALX07 variable region and human PD-1 sequences were input into our internal algorithm, XtalFold^®^. After modeling, the resulting structure was inspected in MOE (Molecular Operating Environment) 2022.02 software and the PyMOL Molecular Graphics System, Version 3.0 Schrödinger, LLC.

### Cloning of humanized ALX07 antibodies

The pcDNA3.4 vector was used to construct separate heavy and light chain expression vectors of humanized ALX07. The heavy and light chain variable region genes of humanized antibodies were designed and synthesized, then inserted into the pcDNA3.4 vector between secretion signaling sequences and constant region genes through In-Fusion cloning. The resulting heavy and light chain expression vectors were confirmed by sequencing and expressed via transfection of vector pair combinations.

### Expression and purification

Antibodies were transiently expressed in suspension ExpiCHO-S cell culture. ExpiCHO-S cells were maintained and transfected in ExpiCHO™ expression medium (Gibco, A2910002) under conditions of 120rpm, 37°C, 8% CO_2_ and humidified incubation. 24 hours prior to transfection, the culture was seeded to 2.5 × 10^6^ cells/mL in a 25 mL culture media, and the cells were adjusted to 6 × 10^6^ cells/mL on the day of transfection. Plasmids encoding the heavy and light chains were combined in a 2:3 ratio, with a total DNA concentration of 0.8-1 μg per mL of culture. The plasmid DNA and ExpiFectamine™ CHO Reagent (Gibco, A29130) were diluted with OptiPRO™ medium (Gibco, 12309019) and mixed before transfection. After 18-22 hours of culture, 150 μL ExpiFectamine™ CHO Enhancer and 6 mL ExpiCHO™ Feed (Gibco, A29130) were added to the culture. After 8-10 days of culture, the media containing the secreted antibody were harvested by centrifugation and clarified using 0.22 μm syringe filters.

Antibodies were purified from the culture supernatants using a 1 mL HiTrap Mabselect SuRe column (Cytiva, 17543801) for affinity chromatography. PBS (pH 7.4) was used as the binding and wash buffer, and 0.1 M sodium citrate (pH 3.5) was used to elute bound antibodies. Eluted antibody samples were neutralized with 1M Tris-HCl (pH 9.0) and buffer-exchanged to PBS using dialysis devices. The purified antibodies were then filtered through a 0.22 μm filter. The concentration of the purified protein was measured with NanoPhotometer (Implen, N60 Touch). The purified antibodies were then characterized by SDS-PAGE and SEC-HPLC (TOSOH, 008541).

### SPR (surface plasmon resonance) analysis

The Biacore 8K+ (Cytiva) was used to characterize the affinity of ALX07 variants to human PD-1 at 25 °C in PBS-P+ (10 mM phosphate buffer (pH 7.4), 137 mM NaCl, 2.7 mM KCl, and 0.005% surfactant P20). Antibodies were captured on a protein A biosensor chip (Biacore, Cytiva), and the extracellular domain of human PD-1 was injected at varying concentrations at a flow rate of 30 μl/min, with a 2-minute association phase and a 15-minute dissociation phase. The sensor surface was regenerated by 10 mM pH 1.5 Glycin-HCl at a flow rate of 10 μl/min. The association (ka, M^-1^s^-1^) and dissociation (kd, s^-1^) constants were calculated using the 1:1 binding model in the Biacore 8K+ evaluation software.

### Humanness Score Evaluation

Humanness scores were evaluated using BioPhi OASis(Prihoda et al. 2022) and AbNatiV(Ramon et al. 2024) algorithms. Both algorithms were downloaded as stand-alone versions and run locally. For BioPhi OASis, the default parameters were used. For AbNatiV, the humanness score was calculated separately for heavy chains and light chains with the VH and VKappa models, respectively. The average score of heavy and light chains was used for each variant.

### AC-SINS (Affinity-capture self-interaction nanoparticle spectroscopy)

The AC-SINS assay was performed to evaluate the propensity of antibodies for self-interaction. Gold nanoparticles (AuNPs, 15705; Ted Pella Inc.) were coated with 80% capturing goat anti-human IgG Fc antibody (109-005-098; Jackson ImmunoResearch) and 20% polyclonal goat non-specific antibody (005-000-003; Jackson ImmunoResearch). After blocking with thiolated PEG (729140, Sigma-Aldrich), the antibody-coated AuNPs were concentrated and incubated with antibodies of interest for 2 hours. The absorbance spectra (510 to 570 nm, 2 nm stepwise) of each well were recorded using a Tecan Spark Multimode microplate reader with SparkControl software. The self-interaction propensity was indicated by a shift in the maximum absorption peak towards longer wavelengths compared to PBS.

### Melting temperature (T_m_) determination

Melting temperature (T_m_) was determined using a QuantStudio Pro6 Real-Time System (Applied Biosystems). Briefly, 39 μL of 0.75 mg/mL antibody was mixed with 1 μL of 200× SYPRO orange dye (S5692, Sigma-Aldrich). The mixture was scanned from 25 °C to 99 °C at a rate of 0.05 °C/min. The melting profile was analyzed using Protein Thermal Shift software, and the T_m_ was assigned from the derivative of the raw data.

### BVP Binding ELISA

The BVP binding ELISA was used to assess the antibody poly-reactivity. Baculovirus particle stock (produced in house) was diluted in 50 mM sodium carbonate (pH 9.6) and coated onto ELISA plates (40303; Beave) overnight at 4 °C. After blocking with blocking buffer (PBS with 0.5% BSA) at room temperature for 1 hour, 100 μL of 0.015 mg/mL testing antibodies in PBS was added and incubated for 1 hour. Bound antibodies were detected using goat anti-human IgG-HRP conjugate (A0170, Sigma-Aldrich) followed by TMB substrate (PR1200; Solarbio). The absorbance was read at 450 nm, and the BVP score was determined by normalizing the absorbance in control wells.

### DNA Binding ELISA

DNA binding ELISA was performed similarly to BVP binding ELISA. DNA (D1626, Sigma-Aldrich, 10 μg/mL in PBS) was coated onto ELISA plates (40303; Beave) overnight at 4 °C. The subsequent steps followed the same procedures as the BVP assay described above.

### Standup monolayer adsorption chromatography (SMAC)

To determine the hydrophobicity of a given mAb using SMAC, 10 μg of sample was injected into a Zenix-C SEC-300 column (213300-4630; Sepax Technologies). A A running buffer containing 150 mM sodium phosphate at pH 7.0 was used with a flow rate of 0.40 mL/min. The retention time for each sample was assigned based on the major peak.

### Cellular binding

HEK293T-HPD1 (CBP74042, Nanjing Kebai) cells were harvested and suspended in FACS buffer (PBS pH 7.2 with 2% Fetal Bovine Serum). Antibodies were diluted in FACS buffer at 8 doses with the final concentration of each dose being 100 nM, 20 nM, 4 nM, 0.8 nM, 0.16 nM, 0.032 nM, 0.0064 nM, and 0.00128 nM. Antibodies were incubated with cells at 4°C for 1 hour. After washing, a secondary antibody (Goat anti-Human IgG (H+L) Cross-Adsorbed Secondary Antibody, Alexa Fluor™ 488, (A-11013, Invitrogen) was added to the stained cells and incubated at 4°C for 30 minutes. Cells were washed, resuspended, and analyzed using iQue3Plus flow cytometry.

### NFAT-luciferase reporter assay

CHO-PDL1 (Nanjing Kebai, CBP74066) cells were seeded in a 96-well plate at a density of 3.5 ×10^4^cells/well and incubated for 16 hours. Jurkat-NFAT-LUC-PD1 (Nanjing Kebai, CBP74018) effector cells were then added at a density of 1.4×10^5^cells/well, along with antibodies diluted in RPMI-1640 medium to final concentrations of 100 nM, 20 nM, 4 nM, 0.8 nM, 0.16 nM, 0.032 nM, 0.0064 nM, and 0.00128 nM. After 4 hours of incubation at 37°C, luciferase activity was measured using a one-step luciferase assay reagent (BPS,78263-5) and detected with a microplate reader (Tecan Spark).

## Notes

### Competing Interest Statement

The authors have declared no competing interest.

