## Supplementary materials for "AI-Guided Precision in Antibody Humanization: Structural Modeling to Minimize Immunogenicity and Preserve Efficacy"

**Supplementary Figure 1. Traditional CDR graft design failed in mab033 humanization.**

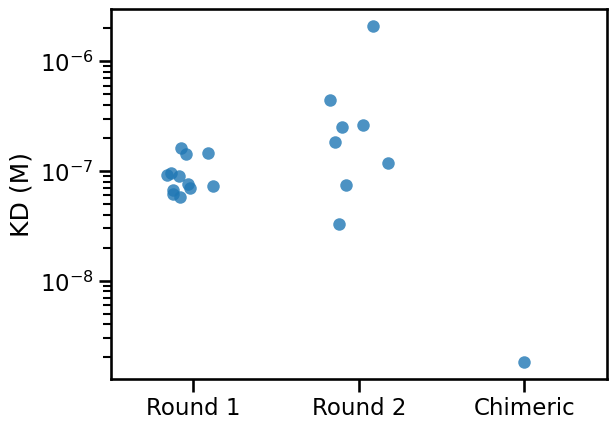

**Supplementary Figure 2 Predicted antibody Fv region structures overlaying on different algorithms.**

Pink: XtalFold^®^, green: ABodyBuilder2, palecyan: IgFold

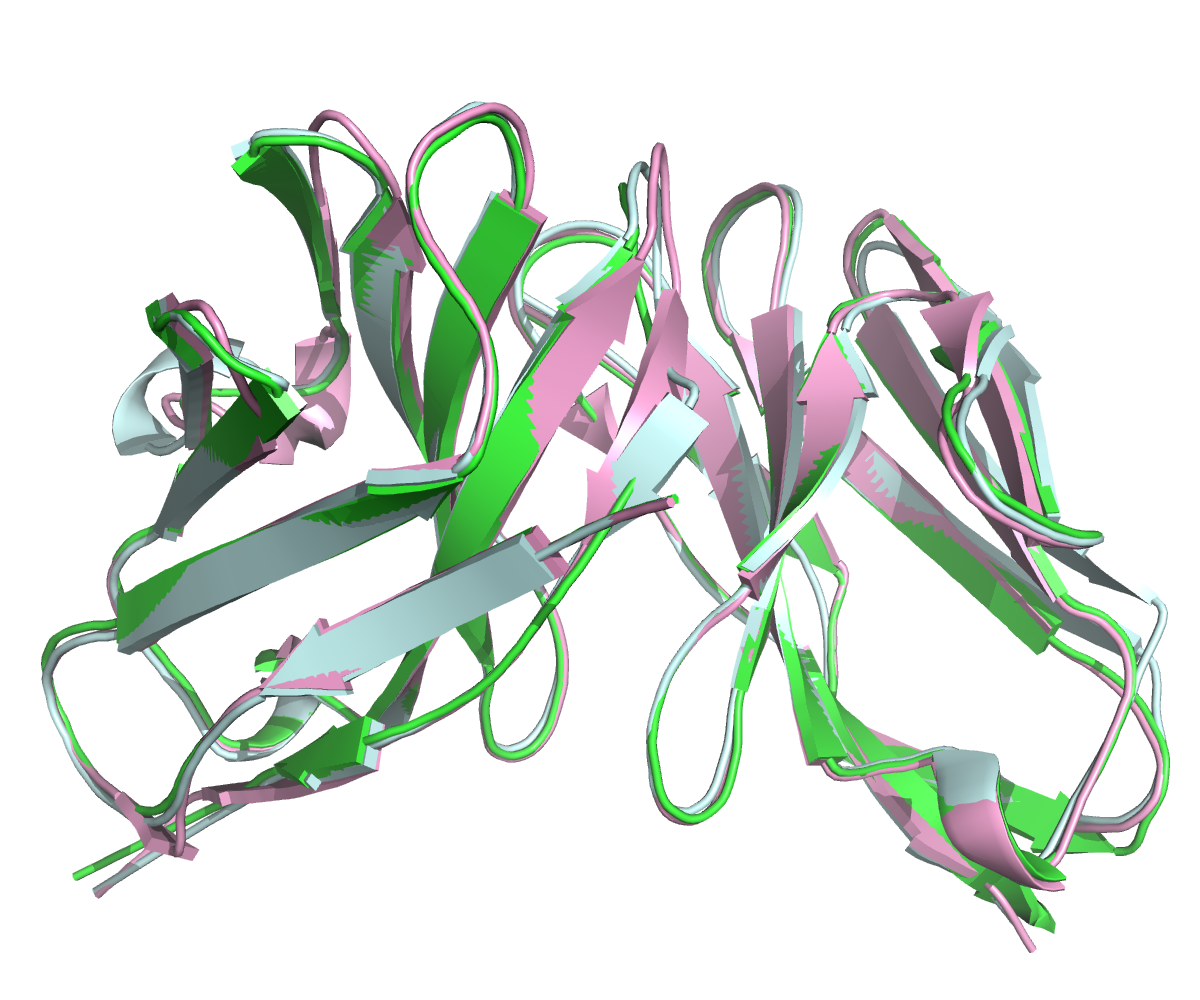

**Supplementary Figure 3. AbNatiV score and OASis percentile score show good correlation, the Pearson correlation coefficient is 1.00**

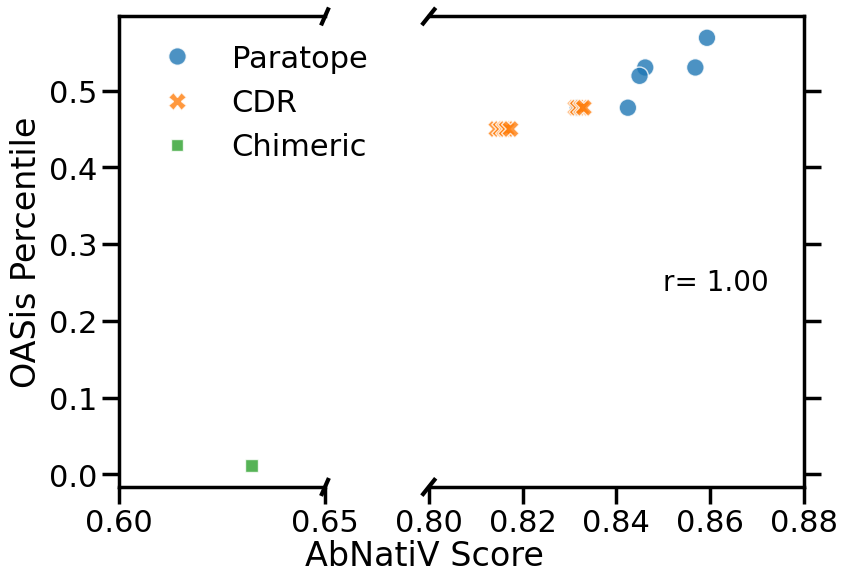

**Supplementary Figure 4. OASis Percentile shows significant differences between paratope and CDR graft designs.**

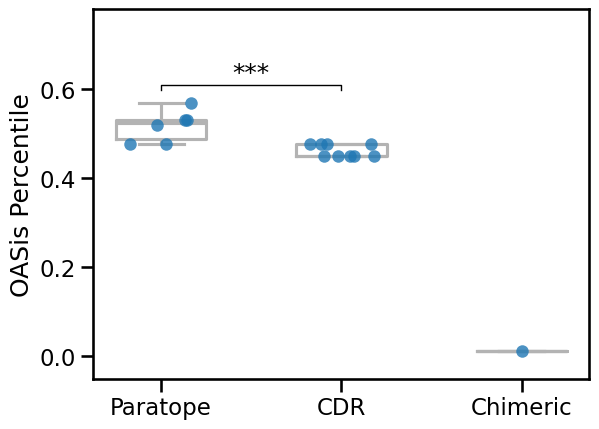

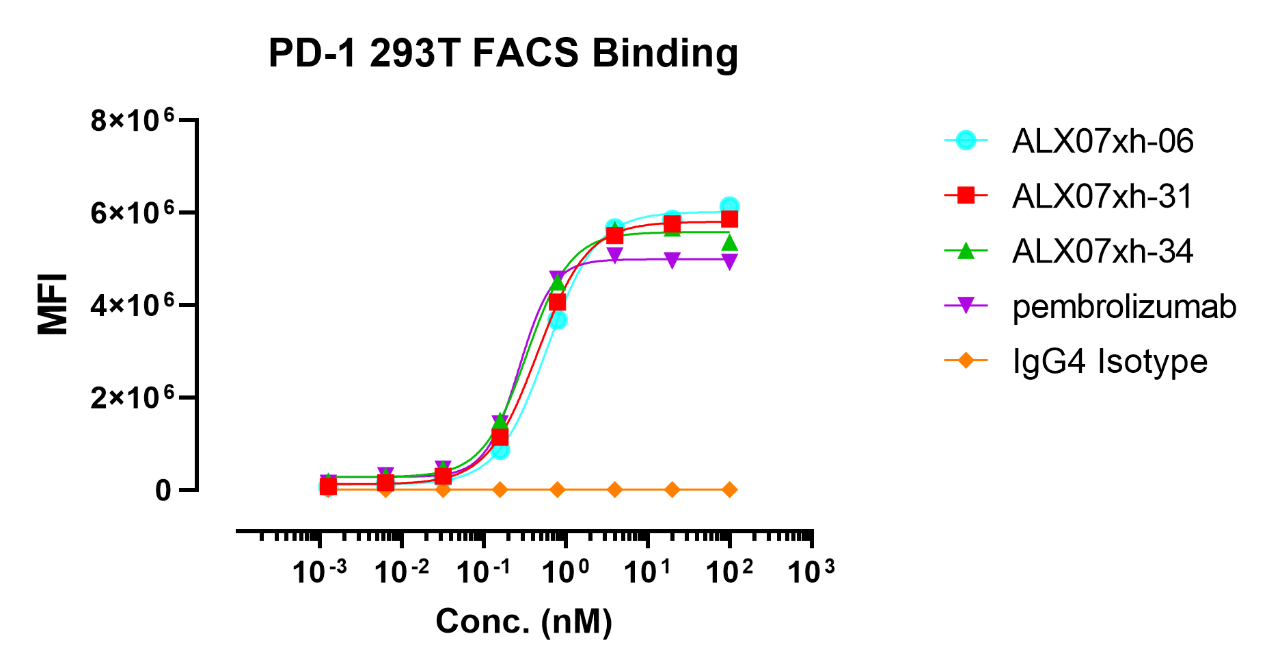

**Supplementary Figure 5A Binding of mab033 humanization variants to the PD-1 over-expression cell line.**

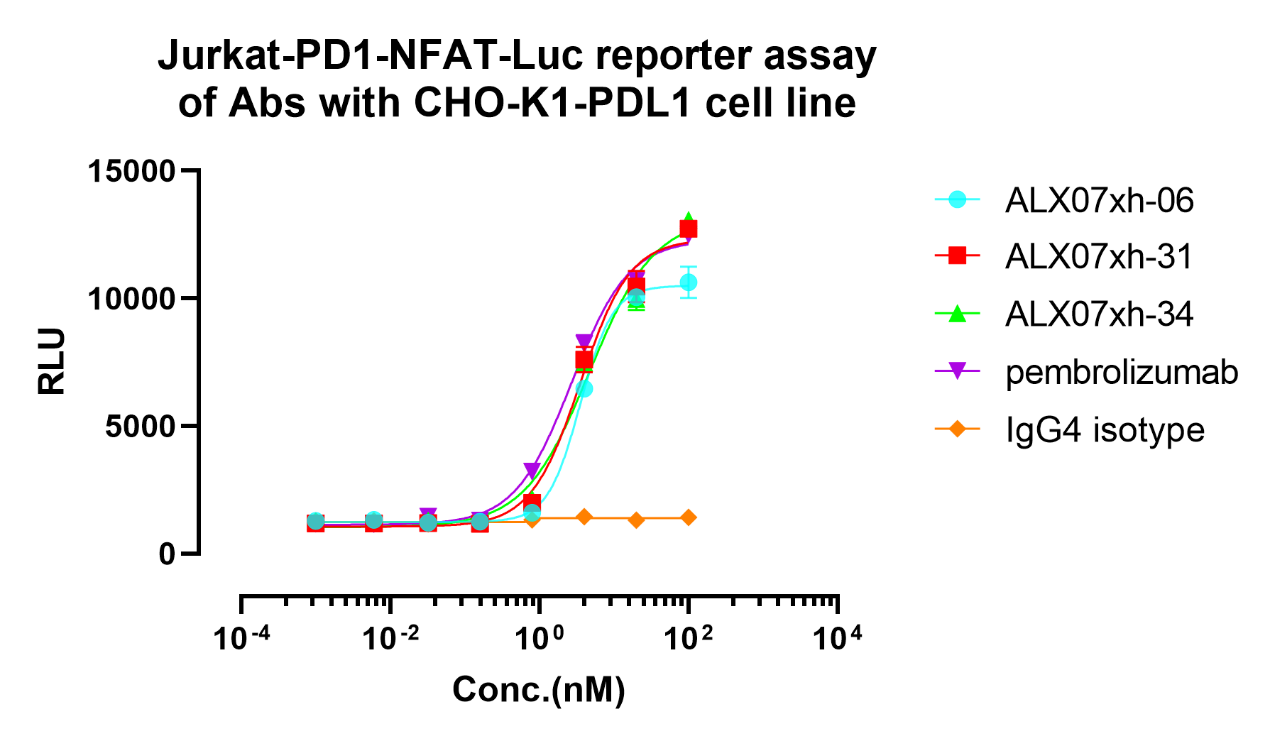

**Supplementary Figure 5B. The reporter assay of mab033 humanization variants.**

**Supplementary Table 1**

Root Mean Square Deviation (RMSD) comparison between XtalFold^®^ predicted antibody structure to ABodyBuilder2 and IgFold.

| Model | HCDR1 | HCDR2 | HCDR3 | LCDR1 | LCDR2 | LCDR3 | FR |
| --- | --- | --- | --- | --- | --- | --- | --- |
| IgFold | 0.68 | 0.82 | 1.63 | 0.61 | 0.78 | 0.83 | 0.49 |
| AntibodyBuilder2 | 0.46 | 0.98 | 2.07 | 0.57 | 0.72 | 0.70 | 0.50 |

**Supplementary Table 2: Paratope-adjacent Residue Differs in Murine and Human V gene**

| position | murine | human | region |
| --- | --- | --- | --- |
| *IGKV4-1*01* | | |  |
| 25 | A | S | CDR1 |
| 30 | S | K | CDR1 |
| 33 | V | L | CDR1 |
| *IGHV2-70*04* | | |  |
| 34 | L | M | CDR1 |
| 35 | G | R | CDR1 |
| 37 | G | S | CDR1 |
| 52 | H | R | CDR2 |
| 54 | W | D | CDR2 |
| 62 | N | S | CDR2 |
| 63 | P | T | CDR2 |
| *IGHV2-5*01* | | | |
| 34 | L | V | CDR1 |
| 52 | H | L | CDR2 |
| 54 | W | Y | CDR2 |
| 62 | N | S | CDR2 |
